# Xyloglucan remodelling defines differential tissue expansion in plants

**DOI:** 10.1101/808964

**Authors:** Silvia Melina Velasquez, Xiaoyuan Guo, Marçal Gallemi, Bibek Aryal, Peter Venhuizen, Elke Barbez, Kai Dünser, Martin Darino, Aleš Pӗnčik, Ondřej Novák, Maria Kalyna, Grégory Mouille, Eva Benkova, Rishikesh Bhalerao, Jozef Mravec, Jürgen Kleine-Vehn

**Author notes:** Equal contributions (Xiaoyuan Guo, Marçal Gallemi, Bibek Aryal). Correspondence and requests for materials should be addressed to J.K.-V.

## Abstract

Size control is a fundamental question in biology, showing incremental complexity in case of plants whose cells possess a rigid cell wall. The phytohormone auxin is a vital growth regulator with central importance for differential growth control. Here we show that growth inducing and repressing auxin conditions correlate with reduced and enhanced complexity of extracellular xyloglucans, respectively. In agreement, genetic interference with xyloglucan complexity distinctly modulates auxin-dependent differential growth rates. Our work proposes that an auxin-dependent, spatially defined effect on xyloglucan structure and its effect on cell wall mechanics specify differential, gravitropic hypocotyl growth.

---

The phytohormone auxin is a central regulator of plant development and has outstanding importance for differential growth control. Despite its significance, we just start to understand the subcellular mechanisms by which auxin reliant growth programs define the size of a cell that is surrounded by a rigid cell wall structure. The increase in auxin concentration correlates with enhanced cellular expansion up to a cell type-dependent concentration threshold, after which auxin eventually interferes with tissue expansion. Accordingly, auxin signalling steers promotion and repression of cell expansion in a concentration- and cell-type-dependent manner (Sauer et al. 2013). These cellular levels of auxin rely on a complex interplay between its metabolism and intercellular transport (Rosquete et al. 2012, Sauer and Kleine-Vehn 2019). On the other hand, tissue specific expression of auxin signalling components and intracellular auxin transport define the cellular sensitivity to auxins (Barbez et al. 2012, Calderón Villalobos et al. 2012, Béziat et al. 2017). Canonical auxin responses take place in the nucleus via auxin binding to its co-receptors TIR1/AFBs and the transcriptional repressor Aux/IAAs (Dharmasiri et al. 2005a, Dharmasiri et al. 2005b). Auxin-induced cellular elongation requires TIR1/AFBs-dependent genomic auxin responses in hypocotyls (Fendrych et al. 2016). In contrast, auxin-triggered repression of root cell expansion utilizes a TIR1/AFBs-dependent, non-genomic pathway (Fendrych et al. 2016, Fendrych et al. 2018). These findings suggest a complex and context specific control of auxin-dependent growth. Auxin-dependent control of cellular expansion is in part manifested by stiffening or loosening of the cell wall (Majda and Robert 2018), but the underlying molecular mechanisms remain long-standing research questions. The plant cell wall is a complex, composite structure comprised mainly of polysaccharides, such as cellulose microfibrils, branched xyloglucans (XyG), arabinoxylans, a diverse pectin matrix, as well as proteoglycans (extensins and arabinogalactan proteins) (Lampugnani et al. 2018). The acid growth theory proposes that auxin-dependent increase in plasma membrane proton pump activity triggers rapid cell wall acidification. The decrease of extracellular pH initiates a cascade of events, including activation of expansins, which dissociate XyG-cellulose networks and consequently promote cell wall loosening (Cosgrove 2014, Dünser and Kleine-Vehn 2015). However, the complexity of the cell wall and also the concentration- and tissue-dependent effects of auxin questions the universal validity of a single universal growth mechanism [e.g. (Calderón Villalobos et al. 2012, Pacheco-Villalobos et al. 2016, Barbez et al. 2017)]. Here we provide evidences that spatial control of XyG structure contributes to differential growth. The extracellular XyG polymer is made of β-1,4-linked D-glucose with functional glycosyl sidechains. In *Arabidopsis thaliana*, 75% of the glucose units are substituted with a xylosyl residue, which can be further substituted (20-30%) with galactose. This galactose moiety can be further decorated with a fucose or an *O-acetyl* group (overview in Fig.S1) (Schultink et al. 2014, Pauly and Keegstra 2016). Several studies have shown that auxin signalling affects various XyG related genes (York et al. 1984, Talbott and Ray 1992, Abel et al. 1994, Xu et al. 1995, Catalá et al. 1997, Catalá et al. 2001, Sánchez et al. 2003, Vissenberg et al. 2005, Osato et al. 2006), suggesting a link between auxin-signalling and XyG-related processes, but the contribution of such a potential interplay to differential growth remains unknown.

## Results and discussion

In order to study auxin-reliant, differential control of growth, we exposed shoots to a gravitropic stimulus, which activates a complex sequence of events ultimately inducing an asymmetric increase of auxin and consequently cellular elongation at the lower side of the shoot (Rakusová et al. 2011). We initially used pea stems as these not only provide material in quantities sufficient for immunoglycan profiling, but also because they are amendable to local auxin manipulation. We longitudinally dissected gravi-stimulated stems and separated the longer (more elongated, convex) and shorter (less elongated; concave) sides (Fig1.A). On these samples, we performed Comprehensive Microarray Polymer Profiling (CoMPP) (Moller et al. 2007, Kračun et al. 2017), using specific antibodies against different cell wall epitopes (Rydahl et al. 2018). We used a calcium chelator cyclohexanediaminetetraacetic acid (CDTA) to extract the calcium-linked pectin matrix and to liberate the soluble cell wall fraction (Kračun et al. 2017). Subsequently, we used 4M NaOH to solubilize hydrogen-bonded hemicelluloses, including XyG, and other more tightly bound cell wall components (Moller et al. 2007). Notably, the monomeric LM15 antibody, generated against non-galactosylated xyloglucan fragments with XXXG motif (Ruprecht et al. 2017), displayed a notable gravity-induced asymmetry when compared to two other hemicelluloses epitopes, such as xylan (LM10 mAb) and mannan (LM21 mAb) (Fig1.A-C), as well as other cell wall epitopes (Fig.S2A-B). This effect was particularly apparent when using NaOH, which is as noted before more suitable to extract XyGs. To assess if this effect relates to auxin-induced differential elongation, we used local application of auxin (lanoline paste) to pea stems, causing the stem to bend (Fig1.D). In agreement with our gravity experiment, asymmetric application of auxin also induced changes in abundances of LM15 epitopes at the site of application (Fig1.E-F). Our unbiased approach suggests that auxin-induced growth correlates with less substituted XyGs, which we subsequently aimed to molecularly characterize in *Arabidopsis thaliana.* We initially visualised potential changes in XyG structure in dark grown hypocotyls of *Arabidopsis*, using the specific CCRC-M1 antibody, which detects fully branched, fucosylated XyG-epitopes (Pattathil et al. 2010). In agreement with the CoMPP analysis in pea, we observed a gravity-induced increase in CCRC-M1 immuno-labelling at the upper/concave side of dark grown hypocotyls in *Arabidopsis* (Fig.1G-H). To independently consolidate an auxin effect on XyG structure, we repressed nuclear auxin signalling by overexpressing PIN-LIKES (PILS) proteins in *Arabidopsis.* PILS proteins are endoplasmic reticulum (ER) localised auxin transport facilitators and repress nuclear abundance and signalling of auxin, presumably by reducing auxin diffusion into the nucleus (Barbez et al. 2012, Béziat et al. 2017, Feraru et al. 2019, Sun et al. 2020). Notably, *PILS5-*dependent repression of auxin signalling correlated with moderate alterations in extracellular monosaccharides, showing slightly increased galactose as well as mildly decreased levels of rhamnose and xylose in dark grown hypocotyls (Fig.S3A), suggesting alterations in the cell wall. To further depict the XyG structure, we used oligosaccharide mass profiling (OLIMP) by MALDI-TOF/MS (Lerouxel et al. 2002) and most prominently observed an increase in fucosylation of XyGs in *PILS5* overexpressing lines when compared to Wt (Fig.S3B), independently confirming that low auxin signalling correlates with complex XyG structures in *Arabiodopsis.*

**Fig.1.**
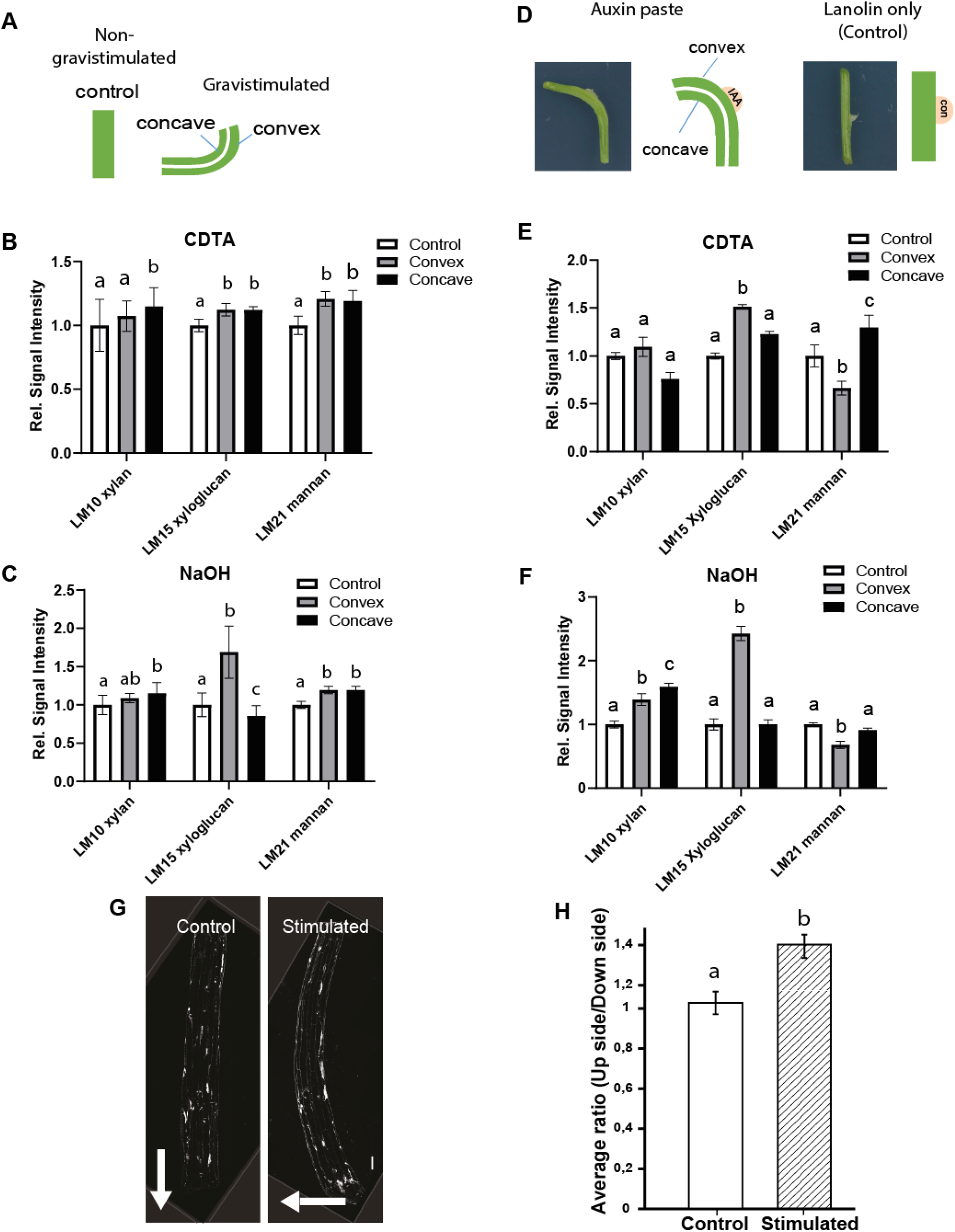
**A-B)** CoMPP profiling of differentially elongated stem segments after gravistimulation. The LM15 antibody, specific to the non-galactosylated (XXXG) motif of XyG, showed increased epitope detection in longer and decreased in shorter stem segments. **A)** Schematic of the experimental design. **B-C)** Quantification of relative changes in the signal intensities with a CDTA **(B)** and NaOH **(C)** extraction in relation to non-stimulated control (n=10, error bars represent SEM). Two-way Anova followed by Tukey with p-value <0.05. **D-F)** CoMPP profiling of pea segments after three-day treatment with an IAA-containing Lanolin paste. **D)** Schematic representation of the auxin application assay. **E-F)** Relative quantifications of both CTDA **(E)** and NaOH **(F)** extractions, in comparison to the non-treated control (n = 7, error bars represent SEM). Two-way Anova followed by Tukey test. **G-H)** Immunolabeling with CCRC-M1 antibody (specific for fucosylated epitopes of XyGs) using longitudinal sections of 5-day dark-grown hypocotyls after gravistimulation and control condition. Two-way Anova with p-value<0.05. **G)** Representative images of sections of non-gravistimulated (Control) and 8hs gravistimulated (Stimulated) dark grown hypocotyls. Scale bar = 100μm. White arrows depict the gravity vector. **H)** Ratio quantification of the mean grey signal at the opposing flanks (e.g. upper and lower) of the hypocotyl section. T-test with p-value<0.05 (n = 4 with 4 technical repeats each).

Using CoMPP, immunocytochemistry, as well as OLIMP approaches, reveal that alterations in auxin signalling correlate with adjustments in XyG structure. Next we aimed to investigated whether repression of nuclear auxin signalling and/or interference with growth correlate with the increase in XyGs structure complexity in dark grown hypocotyls. When grown in darkness, cotyledon-derived auxin stimulates rapid hypocotyl elongation, while suboptimal auxin levels (both the increase and decrease) reduce the expansion of dark grown hypocotyl (Takahashi et al. 2012, Fendrych et al. 2016). To utilize this system for XyG profiling, we next aimed to endogenously decrease and increase auxin levels in dark grown hypocotyls, using estradiol-inducible auxin-conjugating enzyme GH3.6 (Staswick et al. 2005) and auxin-biosynthesis enzyme YUC6 (Cheng et al. 2006). As expected, estradiol-induced overexpression of GH3.6 and YUC6 inhibited hypocotyl expansion, as well as, gravitropic growth (Fig.2A-D).

**Fig.2.**
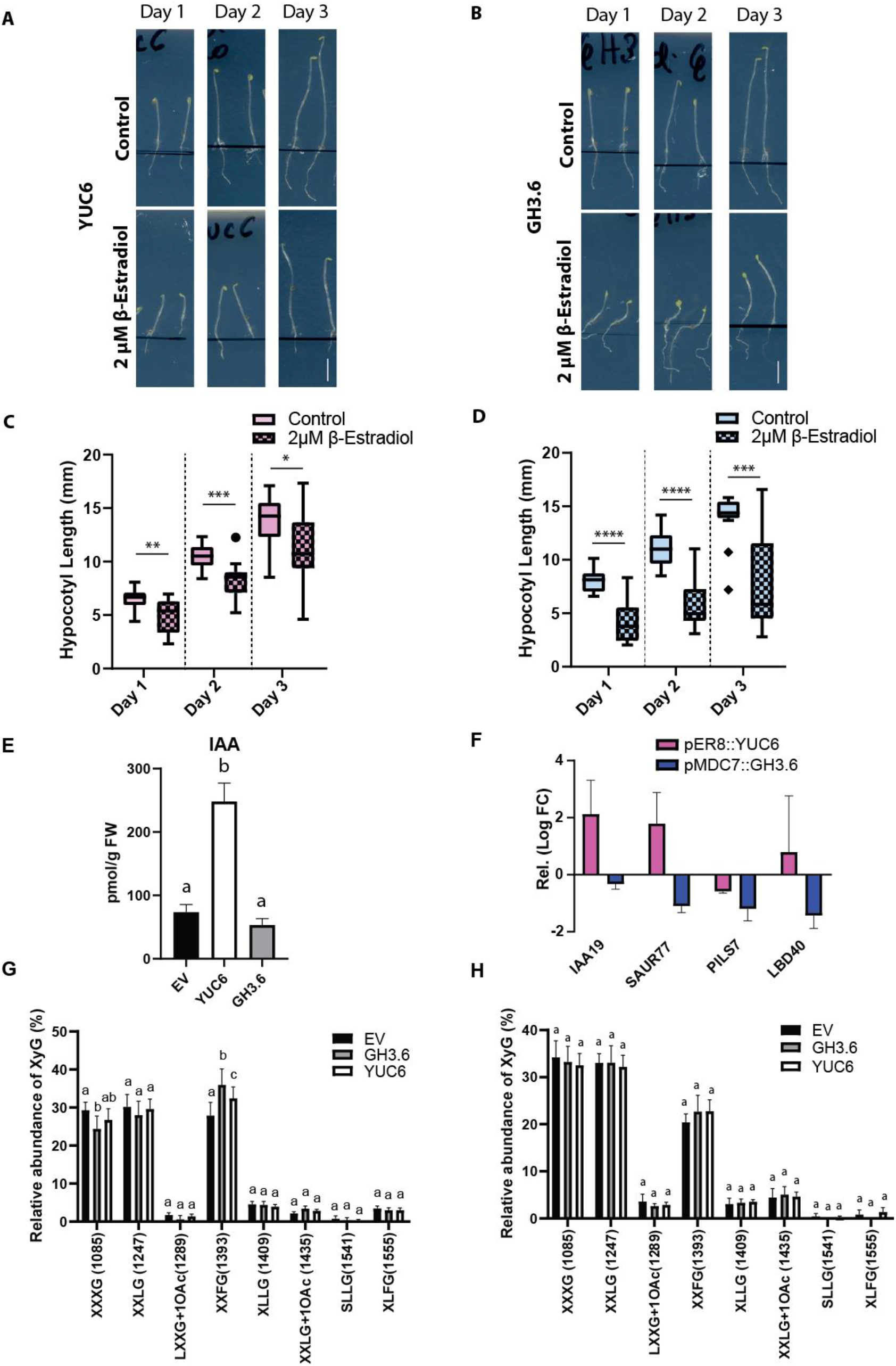
**A-D)** Hypocotyl growth phenotype of estradiol inducible YUC6, GH3.6 and empty vector control (EV). Three-day-old seedlings were transferred onto Estradiol or Control plates and left for 1, 2 and 3 days. **A-B)** Representative images of YUC6 **(A)**, GH3.6 **(B)** on Control media or Estradiol-containing media after 1, 2 or 3 days of induction. Scale bar = 5mm. **C-D)** Quantification of hypocotyl length (Mean±SD) of YUC6 **(C)** and GH3.6 **(D)**. T-test with p-value<0.05. (n = 5-10). **E)** Quantification of free IAA levels of YUC6, G3H.6 and Empty Vector Control (EV). **F)** RT-qPCR of known IAA responsive genes after 3 hours of YUC6 or GH3.6 induction. The Log2 FC values are relative to the treated empty vector control. Representative graph. Data are mean ± SD (n = 3 and three technical repeats for each). **G-H)** OLIMP on upper (growing part) **(G)** and lower (non-growing part) **(H)** sections of hypocotyls of inducible GH3.6, YUC6 and empty vector control (EV). All lines were induced for 3hs with 10μM ß-estradiol. Data are mean± SD (n =4 and 4 technical replicates for each).

We further characterized these lines and detected a YUC6-induced increase of IAA in dark grown hypocotyls (Fig.2E) and only a mild, statistically non-significant reduction in auxin levels after 3h of GH3.6 induction (Fig.3E). Using this condition, YUC6 and GH3.6 induction correlated with increased and decreased expression of known auxin reliant genes, respectively (Fig.2F), confirming its effect on auxin signalling. Therefore, we used this condition to perform an RNA sequencing (RNAseq), intending to fully depict the early auxin reliant transcriptome in these lines. Compared to the empty vector control, 1909 and 2177 genes were differentially expressed after induction of GH3.6 and YUC6, respectively (Dataset S1). The overlapping genes clustered in four categories, displaying (I) up- or (II) down-regulation in both as well as (III) up- and down- or (IV) down- and up- in YUC6 and GH3.6 induced dark grown hypocotyls, respectively (Fig.S4A-B, Dataset S1). Most differentially expressed genes (DEG) (133 and 230 overlapping genes of category I and II) similarly responded to YUC6 and GH3.6 induction (Fig.S4A-B; Dataset S1). On the other hand, we identified 102 genes, which showed inverse (category III and IV) regulation after YUC6 and GH3.6 overexpression. The GO-term analysis (Dataset S2) of these DEGs (Dataset S1) showed as expected enrichment for auxin- and cell wall-related pathways. Among the genes that are structurally related to XyGs, only the galactosyltransferase *MUR3* showed reproducible, but rather mild upregulation after YUC6 induction (Fig.S5A-B).

**Fig.3.**
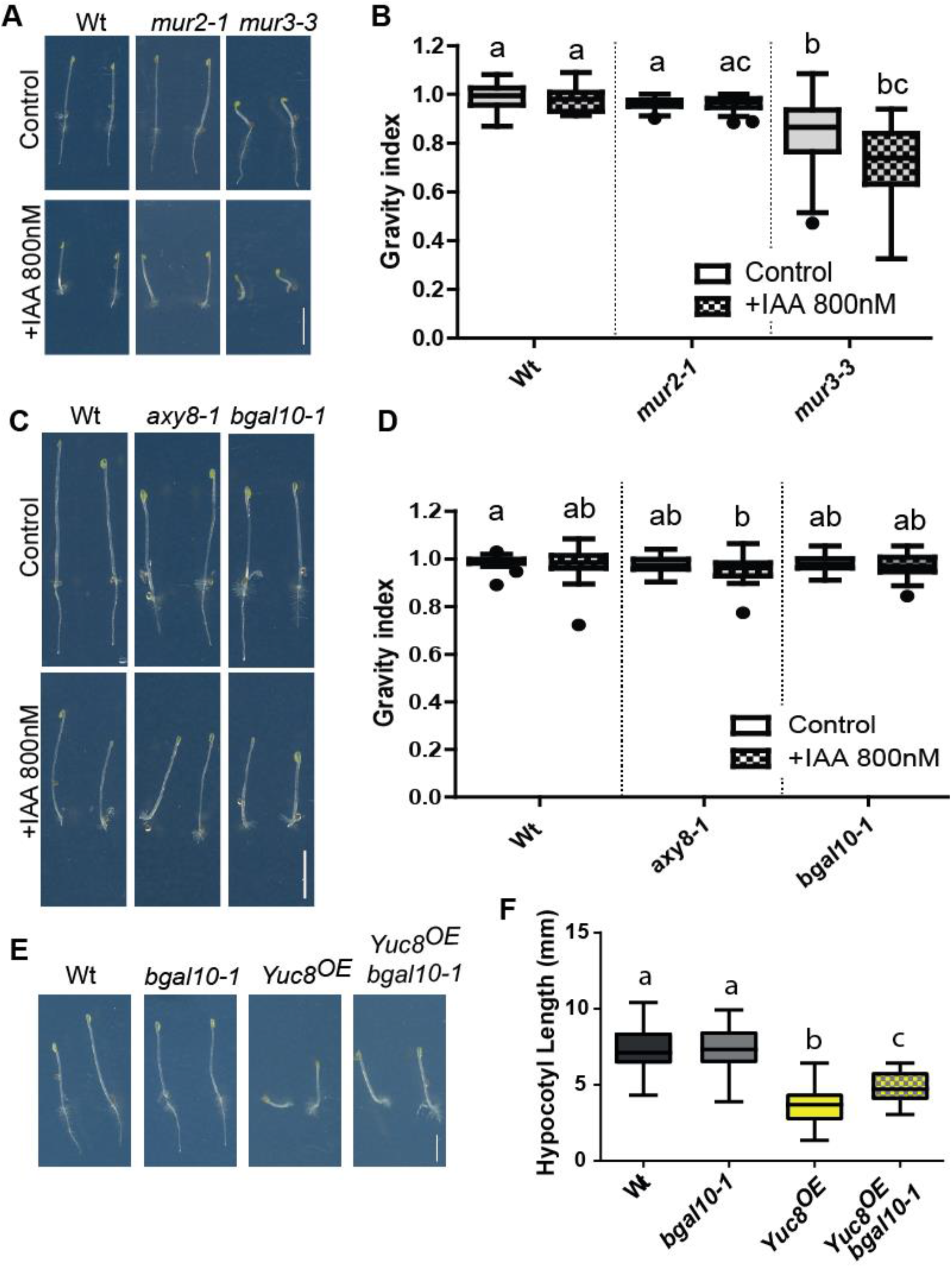
Auxin sensitivity of galactosylation and fucosylation deficient xyloglucan mutants. **A, C)** Mutants of galactosyltransferase *mur3-3* (A), fucosyltransferase *mur2-1* (A), galactosylhydrolase *bgal10-1* (C) and fucosylhydrolase *axy8-1* (C) were grown on 800nM IAA or DMSO (Control) for 5 days. A, C) Representative images of *mur2-*1 (A), *mur3-3* (A), *axy8-1* (C) and *bgal10-1* (C). Scale bar = 5mm. **B, D)** Quantification of the Gravity index of *mur3-3* (B), *mur2-1* (B), *axy8-1* (D) and *bgal10-1* (D) after 800nM IAA or DMSO (solvent control) treatment. This index depicts the ratio between the total hypocotyl length and the shortest distance between the apex and base of the hypocotyl. Tukey box-plot (biological repeats n = 25 – 35). Two-way Anova followed by Bonferroni with p-value<0.05. **E-F)** *bgal10-1* cross with YUCCA8 overexpression line (Yuc8^OE^) **E**) Representative images of dark grown hypocotyls. Scale bar = 5mm. **F)** Quantification of the hypocotyl length. Tukey box-plot (biological repeats n = 35 – 40). One-Way Anova with p-value<0.05.

We next used this system to investigate whether auxin signalling status or the growth context may correlate with complexity of the XyG structures. To address this, we performed OLIMP after YUC6 and GH3.6 induction, which both repress growth but inversely affect auxin levels/signalling. Similar to stable PILS5 overexpression lines, the short-term induction (3 hours) of YUC6 and GH3.6 increased the fucosylation of XyGs in the upper, expanding part of dark grown hypocotyls when compared to empty vector controls (Fig. 2G). Accordingly, we assume that not the auxin signalling levels as such, but the underlying growth response correlates with the detected increase in XyG complexity. In agreement, the lower part of the dark grown hypocotyls, which shows no or reduced expansion rates, did not show the YUC6 and GH3-induced rise in fucosylation of XyGs (Fig. 2H).

Based on these independent approaches, we assume that local auxin concentrations and its underlying growth programs impact on the XyG backbone substitution, where higher and lower XyG complexity may correlate with repression and enhancement of growth, respectively.

We next assessed whether XyGs are required for auxin-reliant differential growth control. Considering the above described alterations in galactosylation and fucosylation of XyGs, we genetically interfered initially with *MUR2-* and *AXY8*-dependent fucosylation and *MUR3-* and *bGAL10*-dependent galactosylation status of XyGs. To assess the overall auxin sensitivity, we exposed the *mur2* and *axy8* as well as *mur3* and *bgal10* dark grown mutants to exogenous auxin (IAA 800nM). Under these conditions, auxin application represses hypocotyl expansion and the XyG related mutants were largely not distinguishable from wild type (Wt) (Fig.3A,C; Fig.S6A-B). On the other hand, we noted that particularly *mur3-3* mutants displayed an exacerbated loss of gravitropic growth when exposed to exogenous auxin (Fig.4A-B). The curved *mur3* dark grown hypocotyl phenotype was reminiscent to seedlings exposed to high levels of auxin (Fig.3A; Fig.S7A). Conversely, hypocotyl expansion of *bgal10* but not *axy8* mutants were partially resistant to high exogenous levels of auxin (Fig.S7A-B) and genetic crosses with *bgal10* mutant partially rescued YUCCA8 overexpression phenotypes (Fig.3E-F; Fig.S7C-D). In order to evaluate the importance of XyGs in auxin-dependent differential growth control, we addressed the gravitropic response in XyG-related mutants. In agreement with presumably hypersensitive auxin responses, *mur3-3* mutant hypocotyls showed gravitropic defects when challenged with a 90° angle change in growth orientation (Fig.4A-B). Compared to *mur3*, gravitropic growth of *mur2-1* mutants were less affected (Fig.4A-B). In agreement with auxin effect on limiting XyG complexity (Fig.1A-H; Fig.S3B), we assume that genetic limitation of XyG complexity correlate with an enhancement of auxin-dependent differential growth. To further asses this, we performed infrared-based growth kinetics of gravitropic dark grown hypocotyls, visualising a hyper bending response of *mur3* mutants (Fig.4C-D), which likely relates to differential cellular elongation rates at the upper and lower hypocotyl flanks (Fig.4E-F).

**Fig.4.**
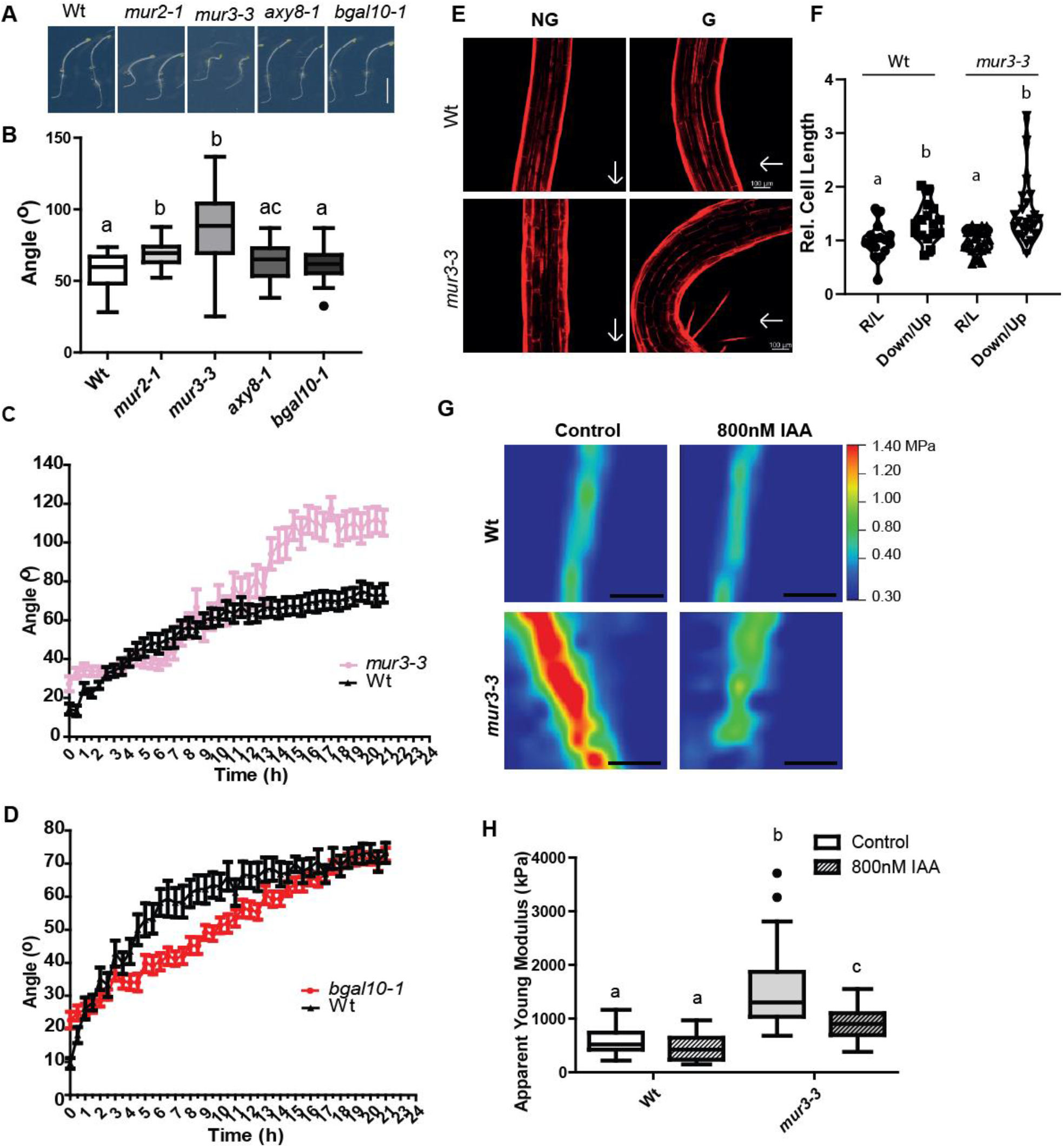
**A-E)** Response to gravistimulation of *mur2-1, mur3-3, axy8-1* and *bgal10-1.* **A-B)** Five-day old dark-grown hypocotyls were challenged with a 90° angle change in growth orientation and the end point angle between the apex of the hypocotyl and gravity vector was measured 24hs later. **A**) Representative images of the angular hypocotyl growth after 24hs. **B)** Quantification of the end point angle. Tukey box-plot. One-Way Anova followed by Tukey. P-value<0.05. Scale bar = 5mm (n = 20-30). **C-D)** Growth kinetics of *mur3-3* **(C)** and *bgal10-1* **(D).** Five-day old dark-grown hypocotyls were challenged with a 90° angle change in growth orientation and placed in a infrared-based dark-imaging box where their growth was recorded. The angle reached every 30 min was quantified. Non-linear fit to a one-phase association curve. K values for each curve were compared. P-value<0.05. Data are mean±SEM (n = 12 – 28). **E-F**) Cell elongation of Wt and *mur3-3* after gravistmulation. Five-day old etiolated hypocotyls were challenged with a 90° angle change in orientation and left overnight, and then stained with Propidium Iodide. **E)** Representative images. Scale bar = 100μm. **F)** Quantification of relative cell length. Between 2-3 cells from the cortex region were measured on the Right (R) side and Left (L) side of Non-Gravistimulated (NG) hypocotyls, or from the Downwards side (Down) or Upwards side (Up) of Gravistimulated (G) hypocotyls. The ratio between R and Left (R/L), and Down and Up (Down/Up) is reported (Mean±SD). Kruskal-Wallis One-Way Anova followed by Dunn test. P-value<0.05. **G-H)** Atomic Force Microscopy (AFM) analysis of *mur3-3.* Dark-grown hypocotyls of *mur3-3* and Wt Col-0 were grown for three days on 800nM IAA or DMSO (Control). **G)** Representative images. Scale bar = 5μm. **H)** Quantification of Apparent Young Modulus in kPa. Tukey box-plot. One-Way Anova followedby tukey. P-value<0.05 (n =30-40).

Conversely to the hyperbending phenotype of *mur3*, the growth kinetics of *bgal10-1* mutants initially showed slower gravitropic bending when compared to Wt (Fig.4D), which also agrees with its partially auxin resistant hypocotyl growth (Fig.3E-F; Fig.S7A-B).

When compared to the fucosylation machinery, the MUR3/bGAL10-dependent galactosylation status of XyGs appears to have a more pronounced developmental importance for auxin-dependent gravitropic hypocotyl growth. We next focused on a family of xylosyltransferases (XXT)1-5, which are responsible for the bulk of the xylosylation of the glucan backbone (Fig.S1). Notably, XyGs were not detectable in *xxt1 xxt2* mutants (Cavalier et al. 2008). Similar to *mur3* mutants, the double mutant *xxt1 xxt2* showed an altered gravitropic index upon auxin application (Fig.S8A-B) and enhanced gravitropic responses (Fig.S8C-D). On the other hand, the phenotype of the *mur3* mutant was fully penetrant and the *xxt1 xxt2* double mutant was under our conditions more variable, which could hint at higher redundancy and/or compensatory mechanisms in this pathway.

Overall, this set of data pinpoint to the importance of XyG complexity for defining auxin-reliant differential growth. Using atomic force microscopy (AFM), tissue mechanics were not altered in shoot apical meristems of *xxt1 xxt2* mutants when compared to Wt, which is likely due to compensatory mechanisms in this tissue (Zhao et al. 2019). On the other hand, a close to 50% reduction in tensile strength was detected for hypocotyls of a partial loss of function allele *mur3-1* (Peña et al. 2004). The time-dependent extension analysis (tensile strength) relates to longitudinal forces along an entire organ and may not strictly correlate with indentation-based (AFM) cellular wall measurements in the epidermis. These two methods may even address the contribution of different cell wall polymers to cell wall parameters (Zhang et al. 2019). To complement the published tensile strength measurements, we performed AFM measurements and, thereby, further assess the contribution of *MUR3* to epidermal cell wall mechanics, using the full knock out mutant *mur3-3.* The epidermal cell walls of untreated dark grown *mur3-3* mutant hypocotyls were much stiffer when compared to Wt (Fig.4G-H), which in principle also correlates with an overall reduction in cell size and hypocotyl growth (Fig.4E-F; Fig.S6A). In combination with previously published work (Peña et al. 2004), this data suggest that *MUR3* contributes to cell wall characteristics in a complex manner. On the other hand, exogenously applied auxin (IAA, 800nM) induced a stronger softening of *mur3-3* mutant cell walls when compared to Wt (Fig.4G-H). Accordingly, *MUR3* seems more sensitive to the auxin impact on wall mechanics, which in principle also agrees with the enhanced gravitropic growth of *mur3-3.* On the other hand, the here observed softening of *mur3* mutant cell walls was not sufficient to induce organ growth (Fig.S6A), suggesting additional growth inhibitory effects when auxin is exogenously applied.

## Concluding Remarks

In conclusion, our data proposes that XyG composition contributes to the rate of auxin-dependent differential growth. We propose that growth inducing and repressing auxin conditions correlate with lower and higher complexity of XyGs, respectively. In agreement, the genetic interference with XyG complexity enhances auxin-reliant differential growth responses. In addition, auxin affects cell wall mechanics in a MUR3-dependent manner. Based on this set of data, we hypothesize that the regulation of XyG complexity and its contribution to cell wall mechanics contribute to auxin-dependent differential growth control. Besides mechanical contributions, the cell wall composition is also sensed and provides a feedback signalling to cellular functions (Vaahtera et al. 2019). In apical hooks, XyG deficiency exerts a negative feedback on the transcription of auxin transport components, abolishing auxin maxima formation and, hence, differential growth (Aryal et al. 2020). In contrast to the apical hook development, auxin-reliant differential growth in gravitropic hypocotyls is not abolished, but on the contrary enhanced in XyG deficient mutants, suggesting a distinct, tissue-specific mode of action. We accordingly propose an unprecedented mechanism assuming that auxin-induced growth signals affect the composition of XyGs, thereby, affecting the wall mechanics for differential growth control in gravitropic hypocotyls.

## Acknowledgements

We are grateful to Paul Knox, Markus Pauly, Malcom O’Neill, and Ignacio Zarra for providing published material; the BOKU-VIBT Imaging Center for access and M. Debreczeny for expertise; and Georg Seifert for critical reading. This work was supported by the Villum Foundation grant 00017489 (JM), Vienna Science and Technology Fund (WWTF) (to J.K.-V.), European Research Council (AuxinER – ERC starting grant 639478 to J.K.-V.), the Austrian Science Fund (FWF) (grant number P26333 to M.K), and the Ministry of Education, Youth and Sports of the Czech Republic through the European Regional Development Fund-Project (CZ.02.1.01/0.0/0.0/16_019/0000827 to A.P. and O.N.).

---

**Fig.S1.**
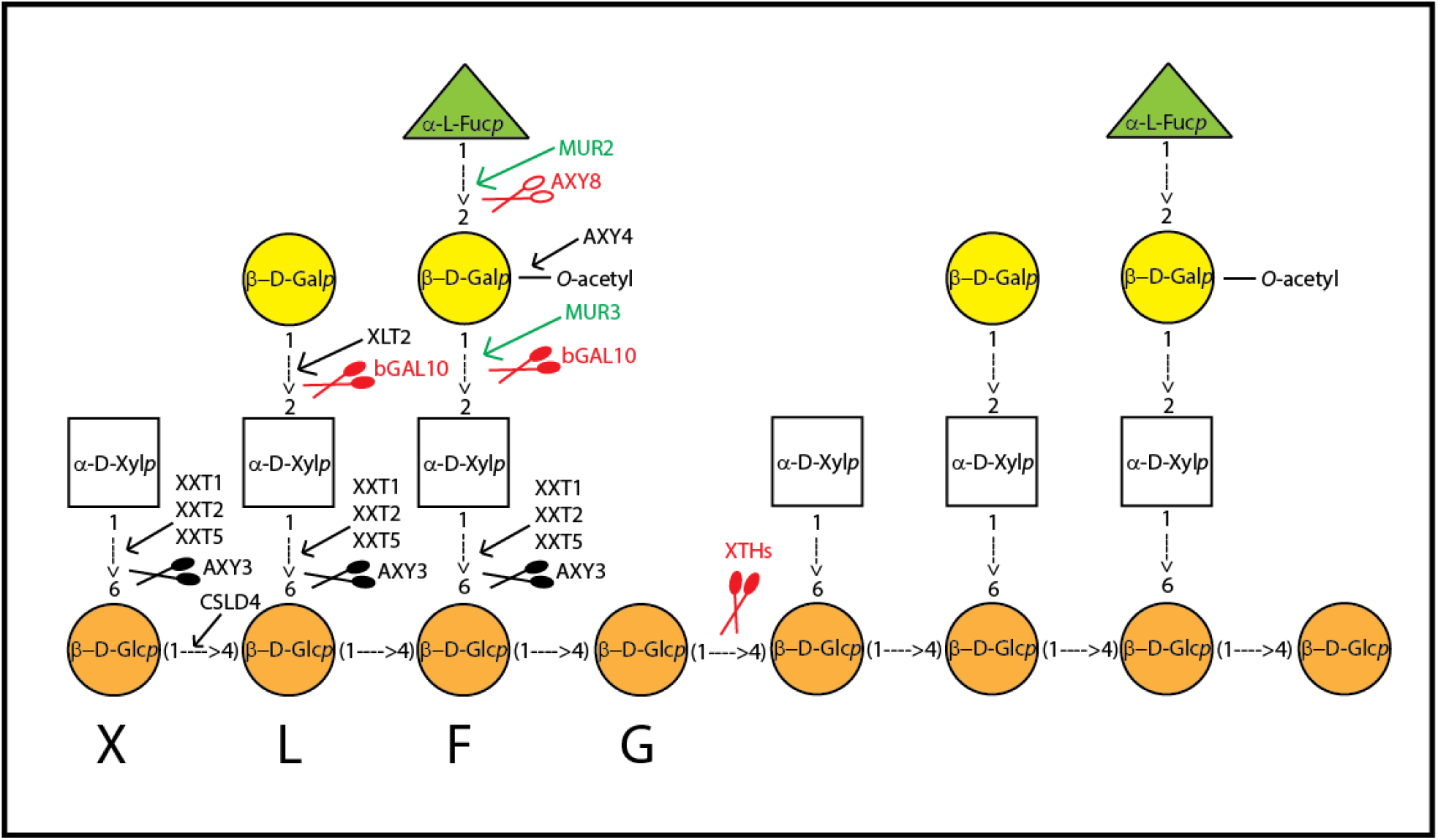
Schematic representation of the biosynthesis and “turnover” of Xyloglucans. The arrows and scissors mark the site of action of the different glycosyltransferases and glycosylhydrolases, respectively. In green, are the biosynthesis enzymes, and in red, the ones that are part of the turnover in the apoplast.

**Fig.S2.**
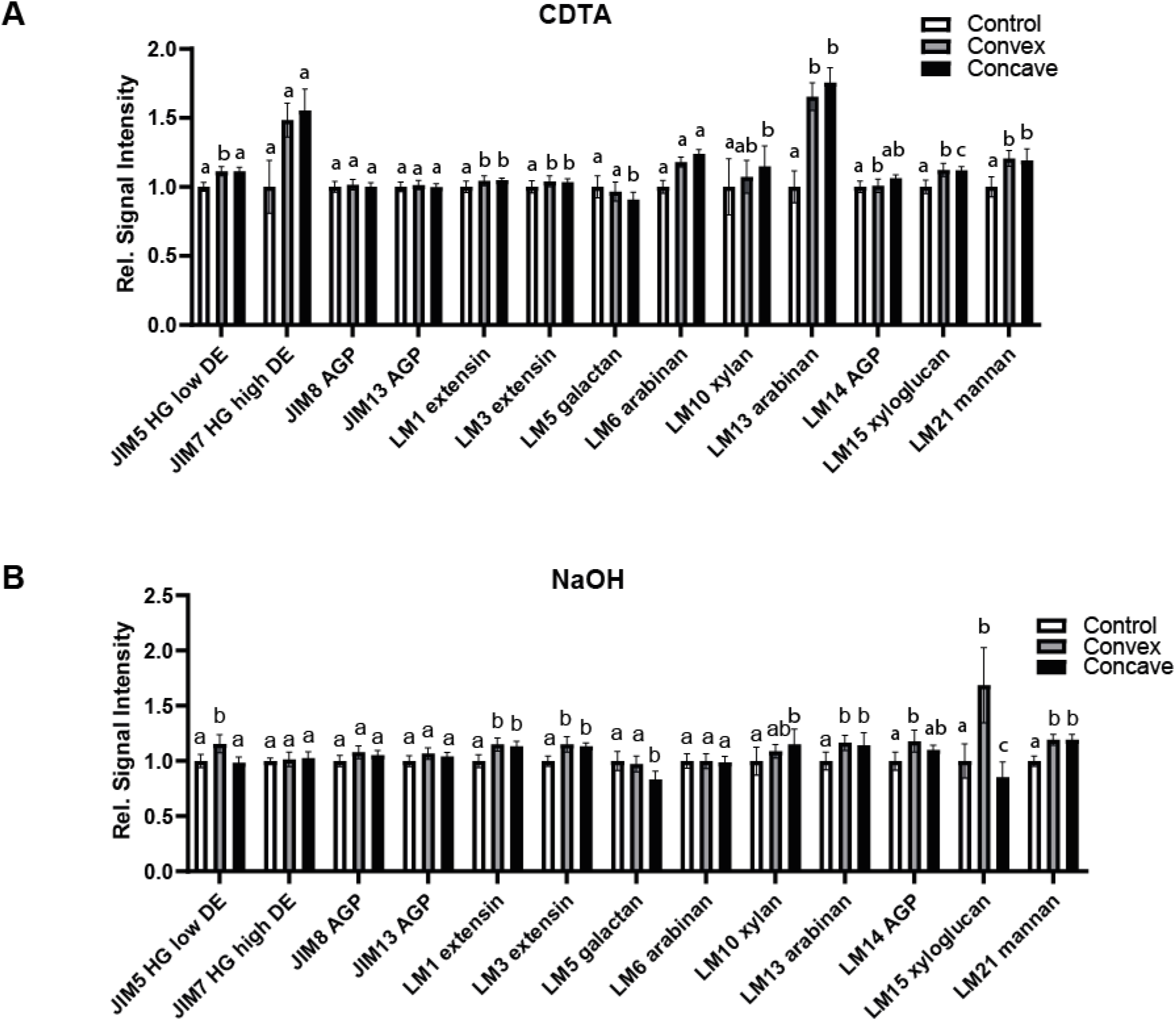
**A-B)** CoMPP profiling of differentially elongated stem segments after gravistimulation. Relative changes in the signal intensities in comparison to non-stimulated control set to 1 (n= 9, error bars represent SEM) with a CDTA (cyclohexanediaminetetraacetic acid) **(A)** and a subsequent NaOH **(B)** extraction. Two-way Anova followed by Tukey with p-value<0.05.

**Fig.S3.**
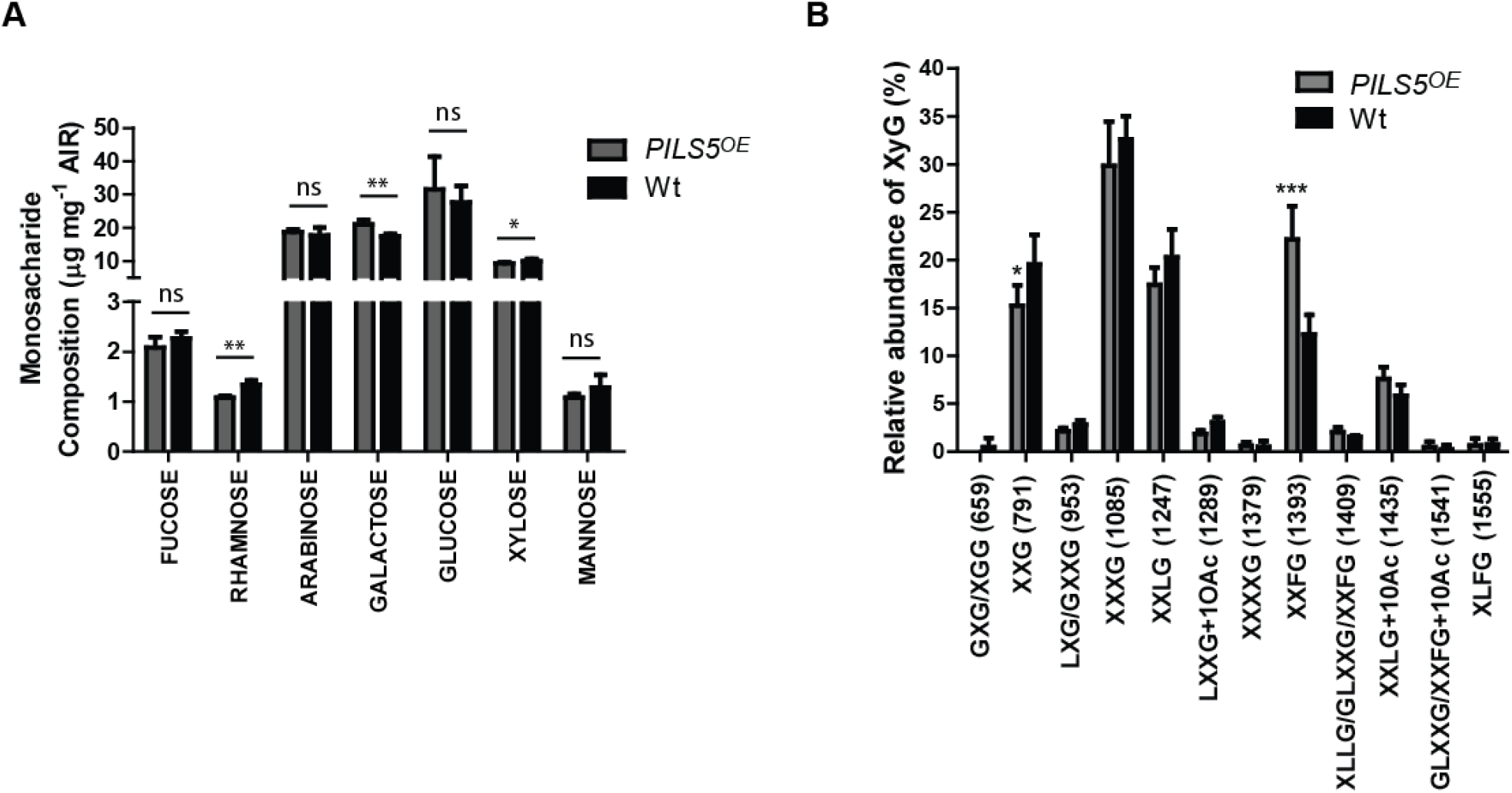
**A)** Monosaccharide composition of wall preparations from etiolated seedlings (n= 4 with 4 technical replicates each). T-test with p<0.05. **B)** Oligosaccharide mass profiling (OLIMP) on 5-day old dark-grown hypocotyls of 35s::PILS5-GFP. Data are mean ± SD (n = 3 and three technical repeats for each). T-test with p-value<0.05.

**Fig.S4.**
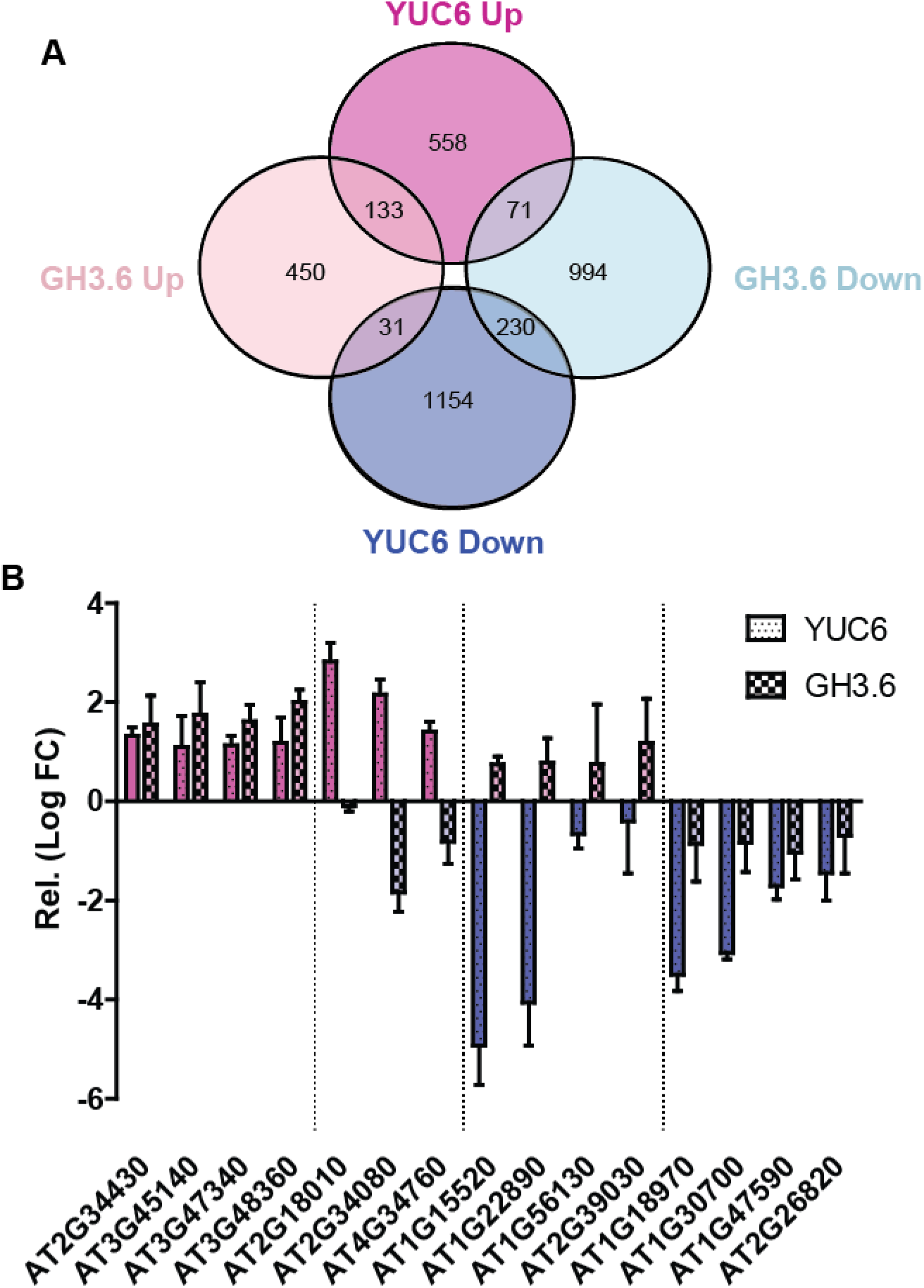
**A-B)** RNA sequencing of estradiol-inducible GH3.6 and YUC6 lines in comparison to empty vector control. Three-day old dark-grown hypocotyls were induced for three hours with 10μM β-Estradiol. **A)** Venn diagram depicting the number of genes that were differentially up-regulated or down-regulated when compared to the estradiol-treated empty vector control. **B)** Validation by RT-qPCR of a subset of the DEG belonging to the four categories in Fig.1A. The Log2 Fold Change (FC) values are relative to the treated empty vector control. Representative graph. Data are mean ± SD (n = 3 and three technical repeats for each).

**Fig.S5.**
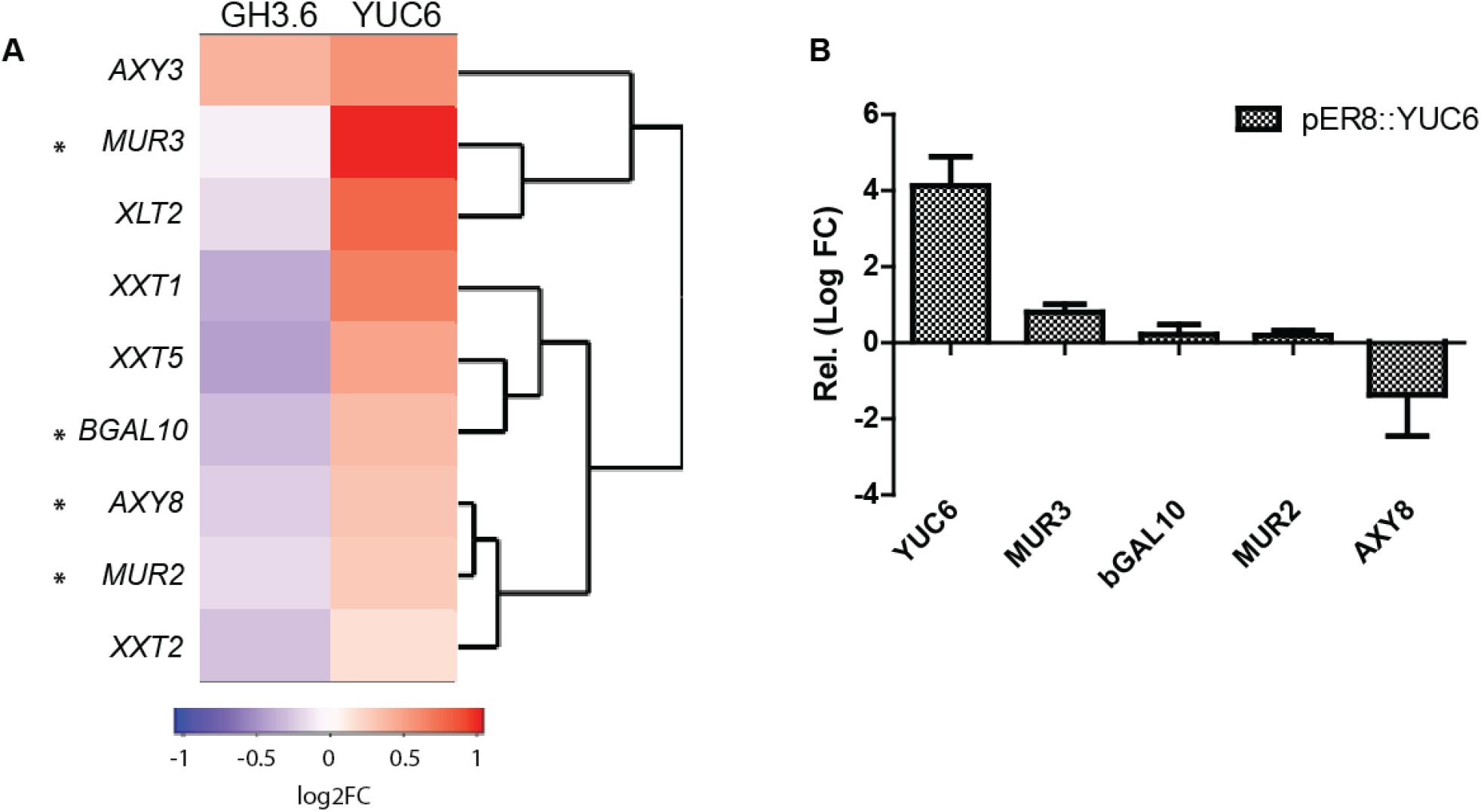
**A)** Hierarchical cluster analysis of gene expression of XyG biosynthesis genes based on log2 FC ratio relative to the empty vector control. Blue represents lowest expression and red highest. Asterisks mark the genes studied in more depth within this work. **B)** RT-qPCR of XyG genes *MUR3, bGAL10, MUR2* and *AXY8* in the estradiol-inducible YUC6. Three-day old dark-grown hypocotyls were induced for 3hs with 10μM β-Estradiol. Induction levels of the *YUC6* gene were evaluated. The Log2 Fold Change (FC) values are relative to the treated empty vector control.

**Fig.S6.**
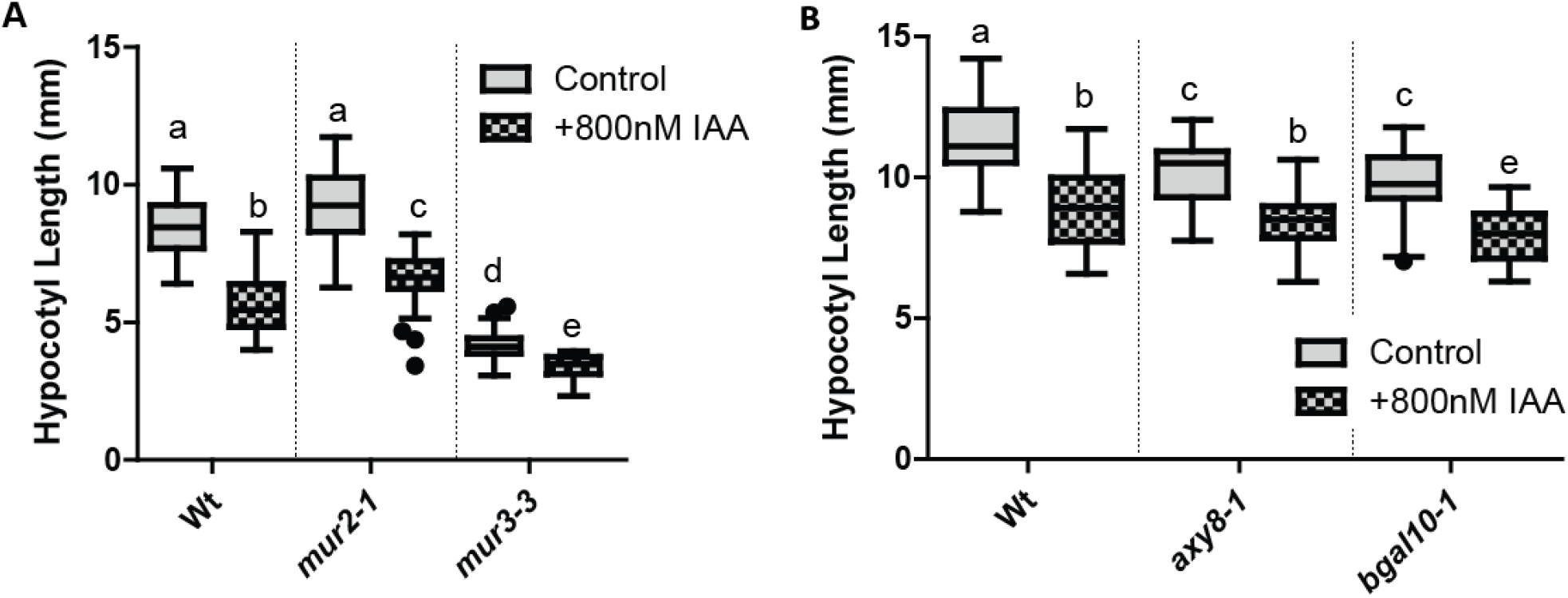
**A-B)** Auxin response of *mur2-1, mur3-3, axy8-1* and *bgal10-1.* All four T-DNA-insertion mutants were grown on 800nM IAA or DMSO (Control) for five days, and then the hypocotyl length was quantified for the treated and control condition. **A)** Quantification for *mur2-1* and *mur3-3.* Tukey box-plot (n = 25 – 35). One-Way Anova followed by Tukey. P-value<0.05. **B)** Quantification for *axy8-1* and *bgal10-1.* Tukey box-plot (n = 25 – 35). One-Way Anova followed by Tukey with p-value<0.05.

**Fig.S7.**
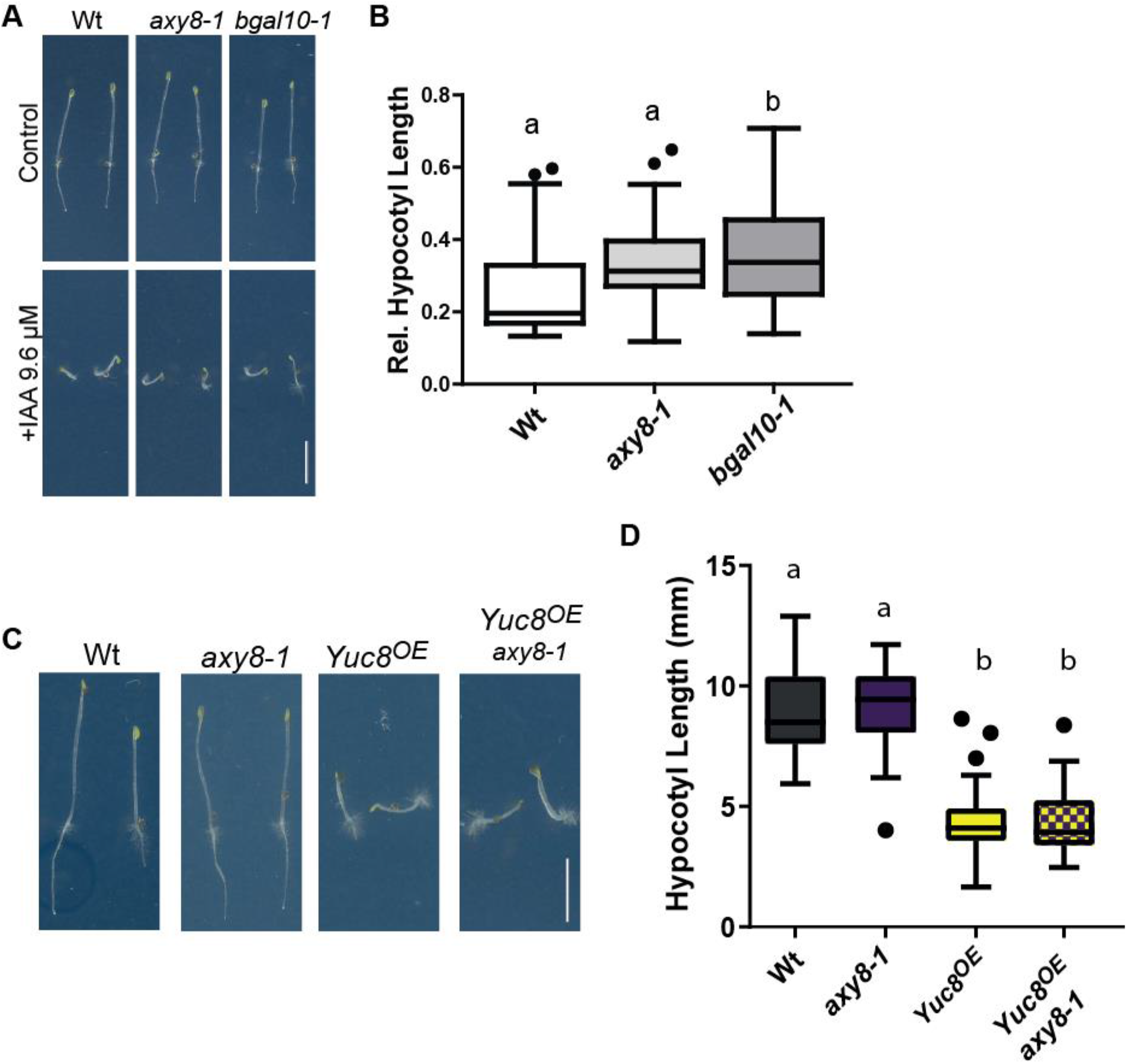
*axy8-1* and *bgal10-1* grown on 9.6 μM IAA or DMSO (Control). **A)** Representative images. Scale bar = 5 mm. **B)** Quantification of the hypocotyl length relative to the DMSO-treated condition. Tukey box-plot (biological repeats n = 35-40). One-Way Anova with p-value<0.05. **C-D)** Genetic interaction between 35s::YUCCA8 *(Yuc8^OE^)* and *axy8-1* crosses. **C)** Representative images of 5-day-old dark-grown hypocotyls of *Yuc8^OE^ axy8-1.* Scale bar =5 mm. **D)** Quantification of the hypocotyl length (Mean±SD) of *Yuc8^OE^ axy8-1.* Tukey box-plot. One-Way Anova followed by Tukey with p-value<0.05. (n = 25-30).

**Fig.S8.**
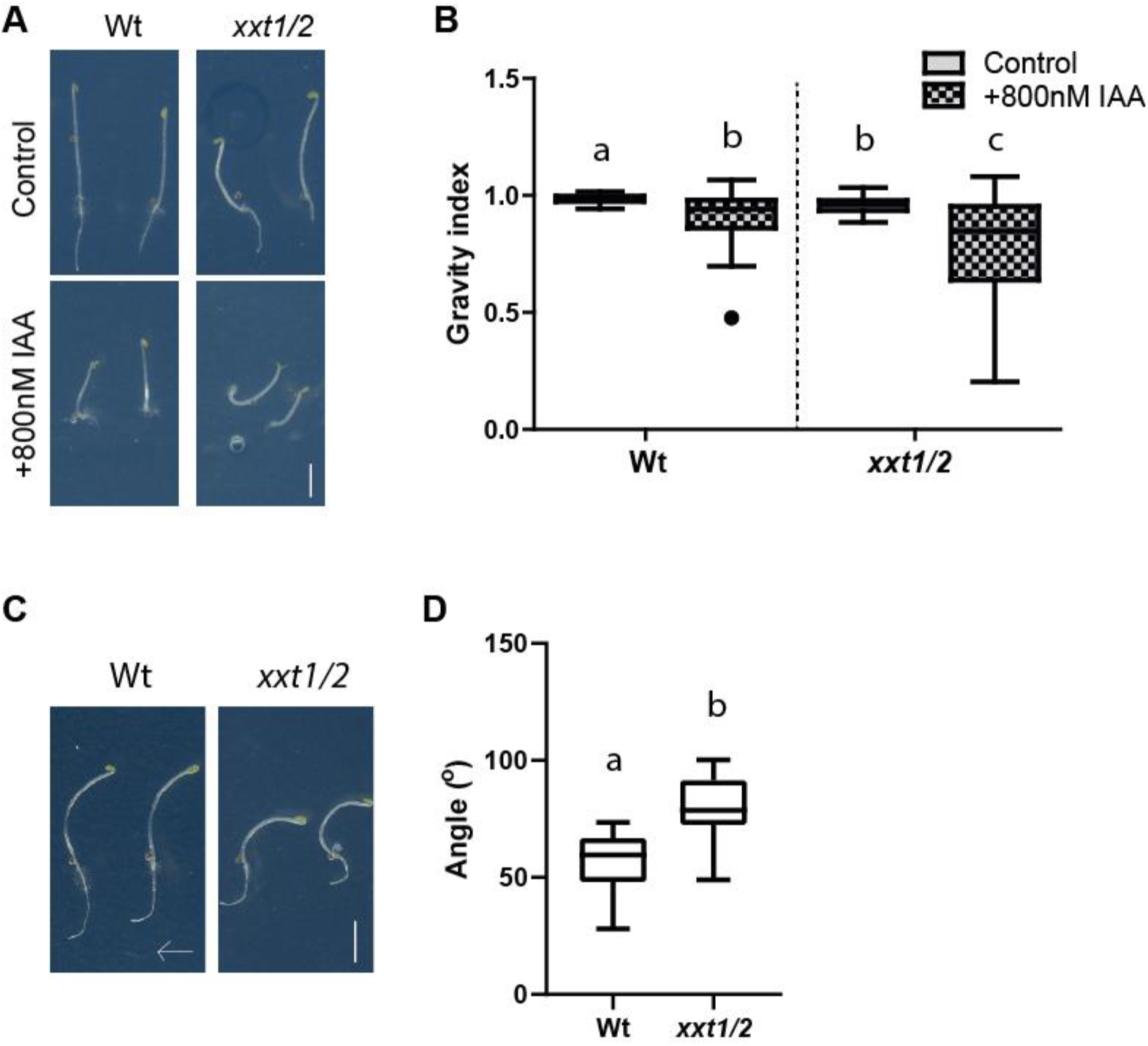
Xylosylation of Xylogulcans. **A-B)** Double mutant of xylosyltransferases *xxt1* and *xxt2* were grown on 800nM IAA or DMSO (Control) for 5 days. **A)** Representative images of *xxt1/2.* Scale bar = 5mm. **B)** Quantification of the Gravity index of *xxt1/2.* One-way Anova followed by Tukey with p-value<0.05. **C-D)** Response to gravistimulation of *xxt1/2.* Five-day old dark-grown hypocotyls were challenged with a 90° angle in growth orientation and the end point angle between the apex of the hypocotyls and gravity vector was measured 24hs later. **C)** Representative images. The arrow marks the gravity vector. **D)** Quantification of the end point angle. Tukey box-plot (biological repeats n = 25 – 27). One-way Anova followed by Tukey with p-value<0.05.

**Dataset S1.** List of differentially expressed genes from the RNA-seq. **SFile1**. Differential expressed genes of pER8::YUC6 vs pMDC7 Empty vector. **SFile2.** Differential expressed genes of pMDC7::GH3.6 vs pMDC7 Empty vector. **SFile3.** Differentially expressed genes shared between pER8::YUC6 and pMDC7::GH3.6 in the up-regulated category. **SFile4.** Differentially expressed genes shared between pER8::YUC6 and pMDC7::GH3.6 that are up-regulated in YUC6 and down-regulated in GH3.6. **SFile5.** Differentially expressed genes shared between pER8::YUC6 and pMDC7::GH3.6 that are down-regulated in YUC6 and up-regulated in GH3.6. **SFile6.** Differentially expressed genes shared between pER8::YUC6 and pMDC7::GH3.6 in the down-regulated category.

**Dataset S2.** GO-term analysis of the four categories determined from the RNA-seq data for the differentially expressed genes in YUC6 and GH3.6 induced dark grown hypocotyls: (I) up- or (II) down-regulation in both, (III) up- and down-regulation or (IV) down- and up-regulation, in YUC6 and GH3.6, respectively. The analysis was performed using the PANTHER Overrepresentation Test.

**Table S1.**
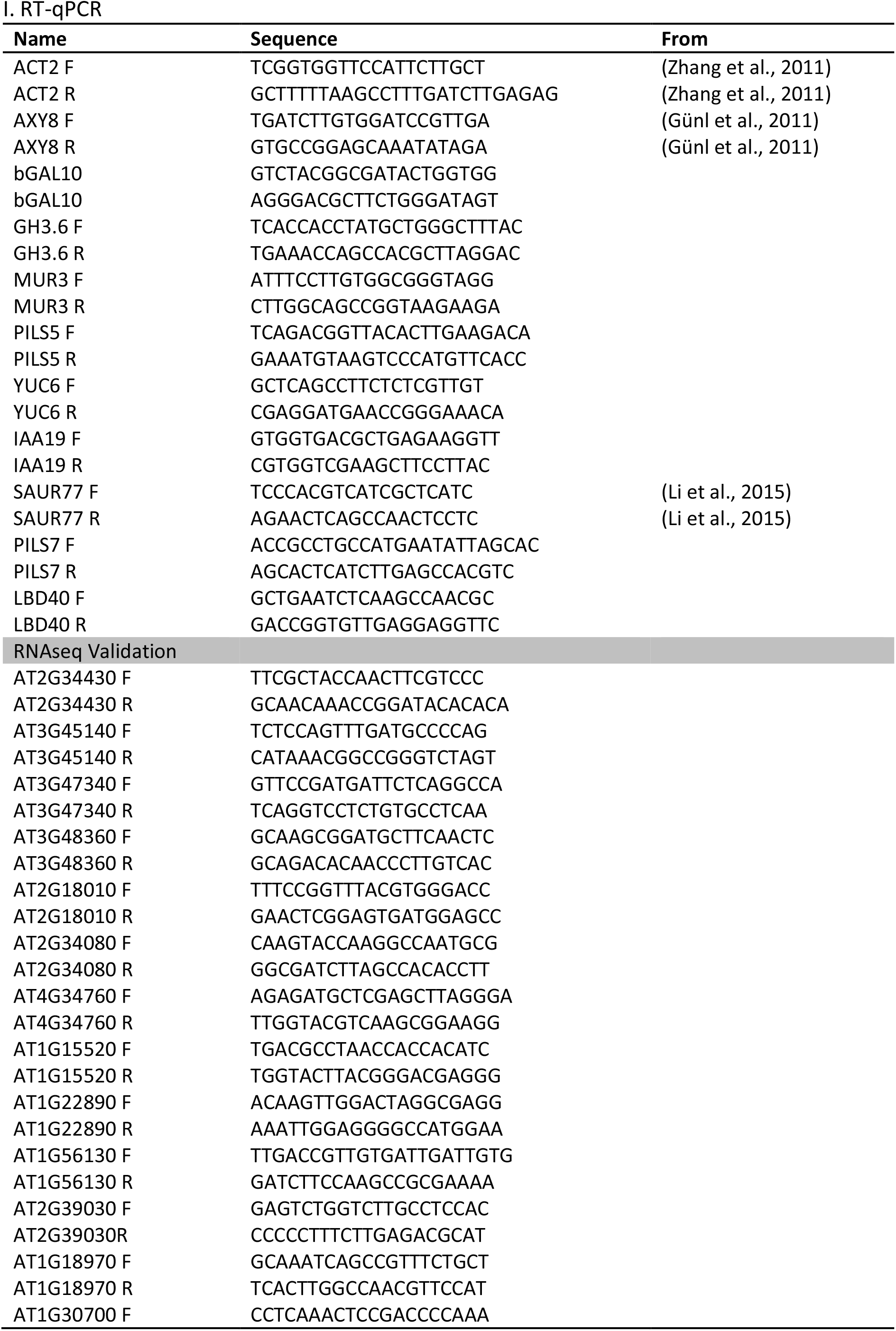
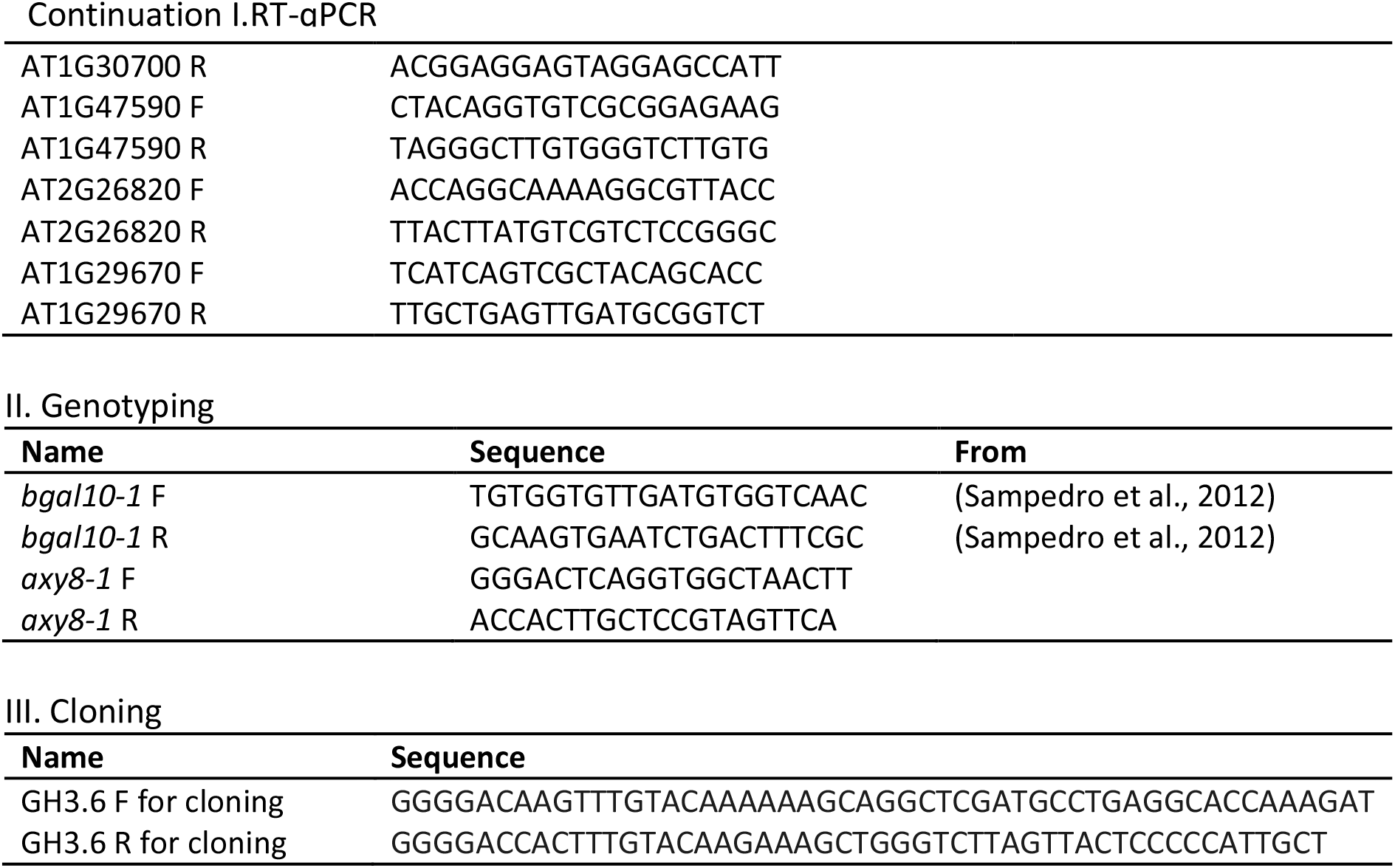
List of primers used in this study.

## Materials and Methods

### Plant material

The Wt background for all lines described is Col-0. Lines *mur2-1* (Vanzin et al. 2002), *axy8-1* (Günl et al. 2011), *mur3-3* (Kong et al. 2015) and *bgal10-1* (Sampedro et al. 2012) have been previously described. The *axy8-1* line was courtesy of Markus Pauly, *mur3-3* was courtesy of Malcom O’Neill and *bgal 10-1* was courtesy of Ignacio Zarra. 35s::PILS5-GFP (PILS5^oe^) was described in Barbez et al. 2012. Primers used for genotyping are listed on Table S1.

### Growth Conditions

Seeds were sterilized overnight with chlorine gas, and afterwards plated in 0.8% agar, 0.5xMurashige and Skoog (MS), and 1% sucrose medium (MS+). For the majority of the experiments (unless stated otherwise), the plates containing the seeds were stratified for two days at 4°C, and after, they were exposed to cool-white light (140μmol.m^-2^.s^-1^) for 8 hs at 21° so as to induce germination, and later kept in the dark for five days at 21°C.

For the auxin treatment experiments, the MS medium was supplemented with 800nM IAA or less than 0.1% DMSO. The seedlings were placed on this medium and grown as described above.

### RNA extraction and RT-qPCR Analysis

We always used hypocotyl tissue for RNA extractions. For the estradiol-induced assays, a 100 μm pore mesh (Mesh Nitex 03-100/44; Transalpina) was placed on top the MS+ medium, and then the seeds were placed on top of this mesh. The plates were then handled as described above for three days (estradiol treatments). At day 3, the plates were uncovered under a green light, so as not to activate any light responses, and the mesh was transferred onto a new plate containing 10μM β-estradiol, and then kept in the dark for 3hs (estradiol), respectively. Tissue was harvested afterwards and total RNA was isolated using the InnuPREP Plant RNA Kit (Analytic Jena), following the manufacturer’s instructions. After RNA extraction, samples were treated with InnuPREP DNase I (Analytic Jena). cDNA was synthesized from 1μg of RNA using the iSCRIPT cDNA synthesis Kit (Bio-Rad) following manufacturer’s recommendations. We used Takyon qPCR Kit for SYBER assay (Eurogentec) and the RT-PCR was carried out in CFX96 Touch Real-Time PCR Detection System (Bio-Rad). ACT2 was used as housekeeping unless stated otherwise. For RNAseq validations, gene AT1G29670 was used as housekeeping, since it was a gene that was stable for all lines and treatments. This gene was selected from the RNAseq data. Primers for all tested genes are listed in STable3.

### Cloning

Gateway cloning was used to construct *pMDC7_B(pUBQ):GH3.6.* The *GH3.6* full-length genomic fragment was amplified by PCR from genomic DNA. Primers are listed in STable3. The PCR was performed using the high-fidelity DNA polymerase “I proof” (Bio-Rad). The full genomic fragments were cloned into the pDONR221 (Invitrogen) vector using Invitrogen BP-clonase according to manufacturer’s instructions. Coding sequences were transferred from the entry clones to gateway-compatible pMDC7_B(pUBQ) vector (Barbez et al. 2012) using the Invitrogen LR clonase according to manufacturer’s instructions. The resulting construct as well as an empty vector were transformed into Col-0 plants by floral dipping in *Agrobacterium tumefaciens* GV3101 strain liquid cultures.

### Quantification of Hypocotyl Length and Gravity index

Seedlings were grown for 5 days in the dark on vertically orientated plates. After this, the plates were scanned with an Epson Perfection V700 scanner. Hypocotyl length was quantified using FIJI 2.0 software (Schindelin et al. 2012).

The Gravity index was calculated as the ratio between the total hypocotyl length and the distance between the apex and the base of the hypocotyl.

For the ß-Estradiol inducible lines pER8::YUC6, pMDC7::GH3.6 and pMDC7 empty vector control (EV), the seeds were first plated onto meshes as already described above and after 3 days, these meshes were transferred onto an MS+ plate containing either DMSO (Control) or 2μM ß-Estradiol, and then left for 1, 2 and 3 days. Afterwards, the plates were scanned and the hypocotyl length was measured as detailed above.

### Gravi-stimulation assays and quantification

Seedlings were grown for four days and then turned 90° and kept in this position for another 24hs. Afterwards, plates were scanned with an Epson Perfection V700 scanner. We measured the angle that was formed between the apex of the hypocotyl and the gravity vector, using the angle tool of the FIJI software.

For the assay where the seedlings were afterwards stained with Propidium Iodide (PI), the gravistimulation was overnight.

### Real time analysis of gravitropic response

Seedlings were grown for 4 days and then turned 90° then placed in this new position in light-sealed box equipped with an infrared light source (880 nm LED) and a spectrum-enhanced camera (EOS035 Canon Rebel T3i) (Béziat et al. 2017). The angles made between the hypocotyl apex and the gravity vector were measured every 30min with the angle tool of FIJI. Representative experiments are shown. Gravitropism kinetics was statistically analyzed using a non-linear regression fit to a one-phase association curve (Béziat et al. 2017).

### Confocal Imaging and Quantification

Imaging was performed using a Leica TCS SP5 confocal microscope, equipped with HyD detector. The fluorescence signal intensity (Mean Gray Value) was quantified using the LEICA LAS AF Lite software. In all cases, a ROI was defined, and the signal intensity was quantified within that region. The same ROI was kept for all analyzed images within said experiment. ROIs used are indicated in the respective figures.

When PI was used, seedlings were incubated for 30min (*mur3-3*), and 1 h (Wt) in a PI solution of 0.02mg/ml.

### RNA-seq

Three-day old seedlings of pMDC7::GH3.6, pER8::YUC6 (Mashiguchi et al. 2011) and pMDC7 empty vector lines were grown and induced as already described above. After the induction time, hypocotyl tissue was harvested and total RNA was extracted using the RNaeasy Plant Mini Kit (Qiagen) following manufacturer’s instructions. Prior to cDNA synthesis, RNA was treated with the RNase-Free DNase Set (Qiagen) with the manufacturer’s recommendations.

The RNA libraries and the subsequent sequencing were performed by the Next Generation Sequencing Facility from the Vienna Biocenter (https://www.viennabiocenter.org/facilities/next-generation-sequencing/). The libraries were generated with the NEBNext Ultra II RNA Library Prep Kit for Illumina with poly(A) enrichment. The sequencing was performed on an Illumina HiSeq2500 with 250bp paired ended fragments.

### Bioinformatics’ Analysis of the RNAseq data

#### Accession numbers

Datasets and NCBI SRA accession numbers will be available shortly.

#### Data pre-processing

Ribosomal RNA reads were removed by mapping the raw reads against the ribosomal transcript sequences using bwa mem [0.7.16a-r1181, (Li 2013)]. The paired end reads were extracted from the unmapped reads using bedtools bamToFastq [v2.29.0, (Quinlan and Hall 2010)] and the Illumina TruSeq adapters were trimmed with cutadapt (Martin 2011).

#### Differential expression analysis

To determine differential expression of the pER8::YUC6 and pMDC7::GH3.6 compared to the pMDC7 Empty vector we considered the transcript per million (TPM) values estimated with Salmon [v0.9.1, (Patro, Duggal et al. 2017)] for the AtRTD2-QUASI transcriptome annotation (Zhang et al. 2017), and used tximport (Soneson et al. 2015) to aggregate the transcript read counts per gene. Differentially expressed genes were obtained with edgeR using the exactTest (Robinson, McCarthy et al. 2010). Genes were considered differentially expressed for a false discovery rate < 0.05.

#### GO-term Analysis

GO-term analysis was performed using the PANTHER Overrepresentation Test (Released 2019.07.11) at http://www.pantherdb.org/. The enrichment was determined comparing the query list of differentially expressed genes with an *A.thaliana* database using a FISHER test with an FDR<0.05.

### Auxin Measurements

Determination of IAA metabolites levels was performed following the methods described before (Pěnčík et al. 2018). As tissue, five-day-old dark-grown hypoctyls of pER8::YUC6, pMDC7::GH3.6 and pMDC7 empty vector control (EV) lines, induced for 3hs on 2 μM β-Estradiol were used. Briefly, 10 mg of tissue were extracted with 1 ml of 50 mM phosphate buffer (pH 7.0) containing 0.1% sodium diethyldithiocarbamate and mixture of stable isotope-labelled auxins standards. 200 μl portion of each extract was acidified with 1M HCl to pH 2.7 and purified by in-tip micro solid phase extraction. After evaporation under reduced pressure, samples were analyzed using HPLC system 1260 Infinity II (Agilent Technologies, USA) equipped with Kinetex C18 column (50 mmx2.1 mm, 1.7 um; Phenomenex). The LC system was linked to 6495 Triple Quad detector (Agilent Technologies, USA).). All samples were measured in quadruplicate for each genotype.

### Atomic Force Measurements and Apparent Young’s Modulus Calculations

The AFM data were collected and analyzed as described elsewhere with minor changes (Peaucelle et al. 2015). To examine extracellular matrix properties, we suppress the turgor pressure by immersion of the seedlings in a hypertonic solution (10% mannitol) for at least 20min before examination. Three-day-old seedlings grown in darkness (in normal AM plate, with or without IAA) were placed in microscopy slides and immobilized using double-glued side tape. We focused on the periclinal cell walls (parallel to growth axis, but perpendicular to the organ surface), and its extracellular matrix. To ensure proper indentations, especially on the regions in the bottom of the doom shape between two adjacent cells, we used cantilevers with long pyramidal tip (14-16 μm of pyramidal height, AppNano ACST-10), with a spring constant of 7.8 N/m. The instrument used was a JPK Nano-Wizard 4.0 and indentations were kept to <10% of cell height (typically indentations of 100-200nm depth and 500nN force). Three scan-maps per sample were taken over an intermediate region of the hypocotyls, using a square area of 25 x 25 μm, with 16 x 16 measurements, resulting in 1792 force-indentation experiments per sample. The lateral deflection of the cantilever was monitored and in case of any abnormal increase the entire data set was not used for analysis. The apparent Young’s modulus (EA) for each force-indentation experiment was calculated using the approach curve (to avoid any adhesion interference) with the JPK Data Processing software (JPK Instruments AG, Germany), based on Herz model adjusted to pyramidal tip geometry. To calculate the average EA for each periclinal wall, the EA was measured over the total length of the extracellular region using masks with Gwyddion 2.45 software (at least 20 points were taken in account). The pixels corresponding to the extracellular matrix were chosen based on the topography map. For topographical reconstructions, the height of each point was determined by the point-of-contact from the force-indentation curve. A total of 12-14 samples were analyzed. A standard t-test was applied to test for differences between genotypes.

### Monosaccharide composition of Polysaccharides

The analysis was performed using four-day-old dark grown hypocotyls on MS half strength supplemented with sucrose. Two grams of this tissue were used to prepare alcohol-insoluble material to be used in the later analysis. For this purpose, hypocotyls were washed twice in four volumes of absolute ethanol for 15 min, then rinsed twice in four volumes of acetone at room temperature for 10min and left to dry under a fume hood overnight at room temperature. For determining the neutral monosaccharide composition, 10 mg of dried alcohol-insoluble material were hydrolyzed in 2.5 M trifluoroacetic acid for 1 h at 100°C as described by (Harholt et al. 2006).

### Xyloglucan Fingerprinting (OLIMP)

Using a green light, four-day old dark-grown seedlings were collected and stored in cold ethanol. Five hypocotyls were dissected for each biological repeat (n = 4), and later used for the analysis. After being left overnight at room temperature in ethanol, the ethanol was removed and the hypocotyls were dried at 37°C for 1 h. Afterwards, 20μl of 50mM acetate buffer, pH5.0, containing endoglucanase from *Trichoderma longibrachiatum* (Magzyme) were added and left overnight at 37°C. OLIMP was then carried out as reported by (Lerouxel et al. 2002) using Super DHB matrix (9:1 mixture of DHB and 2-hydroxy-5-methoxybenzoic acid; Fluka) instead of DHB.

For the pMDC7::GH3.6, pER8::YUC6 and pMDC7 empty vector control (EV), the hypocotyls were grown in the dark for 5 days on top of 100 μm pore meshes MS+ plates and then the meshes were transferred to 2μM β-estradiol MS+ plates. The hypocotyls were dissected into upper section and lower section for the analysis. The rest of the procedure is just as described above.

### Inmunostainings

Arabidopsis hypocotyl section of two days-old seedlings were fixed in 4% paraformaldehyde (PFA) in phosphate buffered saline (PBS) for 45 mins afterward washed 4 times with PBS buffer. Samples were dehydrated for 30 mins sequentially at 30%, 50%, 70%, 90% and 100% EtOH in PBS. LR white was added to samples dropwise to 10% and incubated at 4°C for 6hs. Afterwards, solution was exchanged with 30% LR white in PBS and incubated at 4°C overnight. Solution was exchange with 100% LR white subsequently for 3 times each with 12hs incubation and polymerized at 60°C for 36 hrs. Samples were sectioned at 2.5uM thickness using a Reichert Ultracut S Wild M3Z microtome mounted with a Diatome Histo Diamond Knife (8.0mm 45° angle). Sections were placed on glass slides. Inmunolabelling was performed on sections using CCRC-M1 primary antibody (Agrisera) (Puhlmann et al. 1994) with 1:100 dilution with PBS buffer. Secondary antibody anti Rat Cy5 (Jacksson Immunoresearch) was used with dilution of 1:200. Images were taken using Carl Zeiss LSM780 using 40X magnification (Zeiss C-Apochromat 40x/1.2W Corr M27). Cy5 was excited at 633 nM.

### Geotropism assay

*Pisum sativum* L. cv. Kelvedon Wonder seeds (Frøbutikken, Denmark) were sterilized with hypochlorite solution before germination on wet tissue in sterilised plastic box in dark at room temperature for five days. Seedlings with 3-4cm root were transferred to soil (Pindstrup Mosebrug A/S, Denmark). Plants were grown at 22°C/20°C under 16h/8h day/night temperature cycle in growth chamber under fluorescent light (200 μmol m^-2^s^-1^) for one week. Plants were laid down horizontally for one week. Curvature was harvested and dissected from the middle to get longer (more elongated) and shorter (less elongated) part separately. Stem at the same internode of the plant growing vertically was collected as control. Fresh samples were weighed and homogenized by TissueLyser (Sigma-Aldrich) using a metal bead at max speed. Eight sample pairs were used in the CoMPP study.

### Auxin-induced binding experiment

Pea plants were planted as described above in geotropism experiment. 3-weeks-old pea plant was decapitated until the second internode counting from tip. Lanolin paste containing Indole-3-acetic acid (IAA, Sigma-Aldrich) (10 mg of IAA per 1g of lanoline) was applied to one side of the second internode. Stem applied with plane lanolin paste was set as control. Curvature or straight stems were harvested after 3 days of treatment. The curvature was dissected from middle to separate the longer and shorter side and a pull of three curvatures was used in CoMPP study.

### Comprehensive microarray polymer profiling (CoMPP)

CoMPP analysis was carried out as previously described (Moller et al. 2007). Tissuelysed fresh samples extracted in two solvents sequentially, 50 mM CDTA and 4M NaOH with 1% (v/v) NaBH4 at 30:1 (μl/mg). After 2 hours extraction by shaking and followed 1700 rcf centrifugation, supernatants were printed onto nitrocellulose membrane in four printing replicates and four dilutions in ratio of 1:2, 1:6, 1:18 and 1:54 (v/v). The microarray was probed with a range of cell wall component-directed mAbs (Rydahl et al. 2018) and the intensity of binding was quantified implementing an individual scaling.

## Data Analysis

All graphs and statistical analysis were made with the GraphPad Prismsoftware, versions 5 and 8. Statistical tests are depicted in the figure legends. All experiments were performed at least three times.

